# Study on the interactions between cisplatin and cadherin

**DOI:** 10.1101/2022.09.11.507507

**Authors:** Xitong Wang, Jixiang Sun, Kai Shao, Hongxin Zhi, Yamei Lin, Yujie Fu, Zhiguo Liu

## Abstract

Cisplatin is an important platinum drug in the chemotherapy of many malignant tumors in clinical application. It is well accepted that its main target is DNA molecule. However, recent studies have shown that platinum drugs may also act on some important functional proteins in the human body and have complicate effects on their efficacy. E-cadherin is newly discovered glycoproteins which have been regarded as the important signs of the occurrence and development of some tumors. In this study, the interactions between cisplatin and E-cadherin were explored by fluorescence spectroscopy and atomic force microscopy (AFM). The fluorescence spectroscopy results indicated that cisplatin can efficiently quench the fluorescence of E-cadherin, and the calculated binding constant K_b_ was 3.20×10^6^ (25°C), 1.36 ×10^6^(31°C) and 8.22×10^5^ L mol^-1^ (37°C). The thermodynamic parameters ΔH <0, ΔS <0, ΔG <0, reveal that the fluorescence quenching effect of cisplatin on E-cadherin is static quenching, which is dominated by hydrogen bond and van der Waals interaction. The AFM results revealed that there was a staggered interaction between the intermolecular domains of E-cadherin to form a long spherical chain structure. The addition of cisplatin could significantly affect the staggered effect of the E-cadherin molecules. All these results confirmed that there are strong interactions between cisplatin and cadherin.

## 1 Introduction

No matter in developed or developing countries, the high mortality rate of various types of cancer makes it one of the most important diseases endangering human health. Chemotherapy is one of the main methods in cancer treatment. At present, the widely used anticancer drugs include natural and synthetic alkaloids, taxanes and platinum anticancer complexes. Among them, platinum complexes play an important role in the treatment of various cancers. It is estimated that about 50% of patients use platinum drugs in the chemotherapy of various cancers [1]. The annual sales of platinum anticancer drugs in the world has now reached the order of $2 billion [2]. Platinum anticancer drugs were first discovered by American scientists Rosenberg and others in the 1960s. Rosenberg et al. observed that *E. coli* changed from sausage like to filament like structure after a period of current was applied to *E. coli* in ammonium chloride buffer with a platinum electrode, which was due to the inhibition of its cell division [2]. Subsequently, it was found that this phenomenon was mainly caused by the hydrolysate produced on the platinum electrode. In 1969, Rosenberg et al finally discovered that cisplatin (CIS dichlorodiaminoplatinum, cisplatin, see structural formula 1) has strong anticancer activity [3]. This important discovery opens up a new field of anticancer research of metal complexes. Based on the structure-activity relationship of cisplatin against cancer, scientists try to synthesize new high-efficiency and low toxicity anticancer metal complexes.

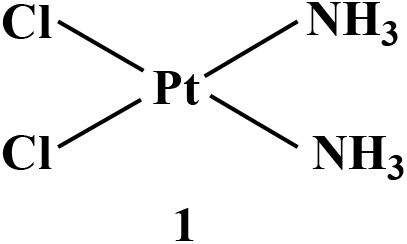
Chemical structure of cisplatin (CIS dichlorodiaminoplatinum)

Recent studies have shown that many small drug molecules can interact with proteins in human body and ultimately affect their pharmacological activities [4]. The main target of cisplatin is DNA molecule. However, it has been found that cisplatin can interact with some proteins to affect their final activity [1]. For example, cisplatin can interact with ubiquitin proteins and can be affected by coexisting glutathione [5]. Cisplatin DNA complex can interact specifically with high mobility protein (HMG domain protein). This specific interaction can have a significant impact on the recognition, clearance and repair of platinum drugs [6].

Recently, researchers found that a large class of transmembrane glycoproteins named E-cadherin are important signs of the occurrence and development of some cancers [7-9]. Cadherins are a kind of calcium ion dependent adhesion glycoproteins. Each cadherin molecule plays an important role in cell-cell adhesion after binding 12 Ca^2 +^. The research report published in Nature confirmed that E-cadherin has an important relationship with the occurrence and development of cancer [8, 9]. In addition, recent studies have shown that the increased concentration of E-cadherin in serum is closely related to tumorigenesis, invasion and metastasis. In particular, the concentration of soluble cadherin in serum was found to be significantly higher in patients with gastric cancer than in healthy controls, which may be due to the degradation of E-cadherin on the surface of tumor cells and its release into the blood. Therefore, the concentration of soluble cadherin in serum can be used as an important indicator for the prevention, diagnosis and treatment of gastric cancer [10, 11].

Cisplatin can interact with human serum albumin in blood, which can significantly influence its pharmacological activity[12]. This result implied that cisplatin may interacted with other proteins in serum such as E-cadherin. To the best our knowledge, there is no systematic study and report on the interactions of cisplatin and cadherin. In this study, we used fluorescence spectroscopy and AFM to analyze the interactions between cisplatin and cadherin.

## 2 Experimental

### 2.1 Materials and instruments

Cisplatin (CIS dichlorodiaminoplatinum (II), cisplatin) was purchased from Kunming Guiyan Pharmaceutical Co., Ltd. E-cadherin human (recombinant) was purchased from sigma Aldrich (Shanghai) Co., Ltd. The original concentration of cadherin was 0.5mg/ml, and the concentration used in the experiment was diluted with phosphate buffer solution. The highly oriented pyrolytic graphite (HOPG) used as the substrate in atomic force microscope observation is the product of Mikro Masch company in Estonia. The pure water used in the experiment was purified by milli-Q system, and the resistivity reached more than 18m.Ω. The fluorescence spectrophotometer equipped with the temperature-controlled water bath device to measure the fluorescence spectrum at different temperatures is F-7000 produced by Hitachi, Japan. The atomic force microscope instrument used in the experiment is molecular imaging Picoplus II system (molecular imaging, USA). The specification of imaging scanning head is a one hundred μm type, which was corrected by standard sample before use. The probe used for scanning is manufactured by Mikromasch, model NSC35, the radius of curvature of the AFM tip is less than 10 nm, the force constant is 7.5n/m, the scanning speed is less than 1lines/s, the resonance frequency of imaging is 130 kHz, and the scanning is carried out in tapping mode under room temperature conditions.

### 2.2 Experimental methods

#### 2.2.1 Fluorescence spectrometric sample preparation

3mg of pure cisplatin powder was weighed, and dissolved in 10 ml of ultrapure water. The cisplatin storage solution has a concentration of 1mmol/L, and store it at 4 °C before use. This 1 mmol / L cisplatin storage solution was diluted with ultrapure water for three gradients to a concentration of 1 μmol/L. 10 μL of original cadherin (0.5mg/ml) solution was put into a 1ml centrifuge tube, and added 410 μL phosphoric acid buffer solution (0.2 mol / L, pH = 7.4) to obtain a concentration of 0.2 μmol / L dilute solution of cadherin. In this cadherin solution, 1 μmol/L cisplatin solution was gradually added, keeping the centrifuge tube in a water bath with the predefined temperature for 5 min at the corresponding temperature. The solution was then transferred into a micro quartz cuvette, and measured by fluorescence spectrophotometer excitation at wavelength of 280 nm, recording its fluorescence spectrum at the range of 290-500 nm.

#### 2.2.2 preparation and characterization of cisplatin-cadherin samples

0.2 μM cadherin solution interacting with cisplatin was diluted ten times with 0.2M phosphate buffer solution and used to prepare atomic force microscope samples. The final concentration of cadherin was 0.02 μM for preparing AFM sample. 50 μL of the final solution was added dropwise to the newly cleaved high orientation pyrolytic graphite (HOPG) surface. The droplets removed on the HOPG after incubation for 10 min, and the surface are soaked in deionized water for three minutes, and then gently rinsed with ultrapure water more than 10 times to remove the loosely adsorbed cadherin. Finally, the samples are dried with inert argon gas, and stored in a desiccator before imaging with an atomic force microscope.

## 3. Results and discussion

### 3.1 Fluorescence quenching of cadherin by cisplatin

Fig. 1 shows the effect of cisplatin, a typical platinum drug, on the fluorescence spectrum of cadherin under different temperature conditions (25 °C, 31 °C and 37 °C). After the excitation of cadherin at 280 nm, the maximum fluorescence emission peak appeared at 338.6 nm. This fluorescence can be attributed to the tryptophan residues in the cadherin structure. With the gradual addition of cisplatin solution, the intensity of its maximum fluorescence emission peak gradually decreased, as shown in Fig. 1 (a) (b) (c). The results show that cisplatin can effectively quench the fluorescence emission of cadherin, indicating that there is a strong interaction between them.

**Fig. 1.**
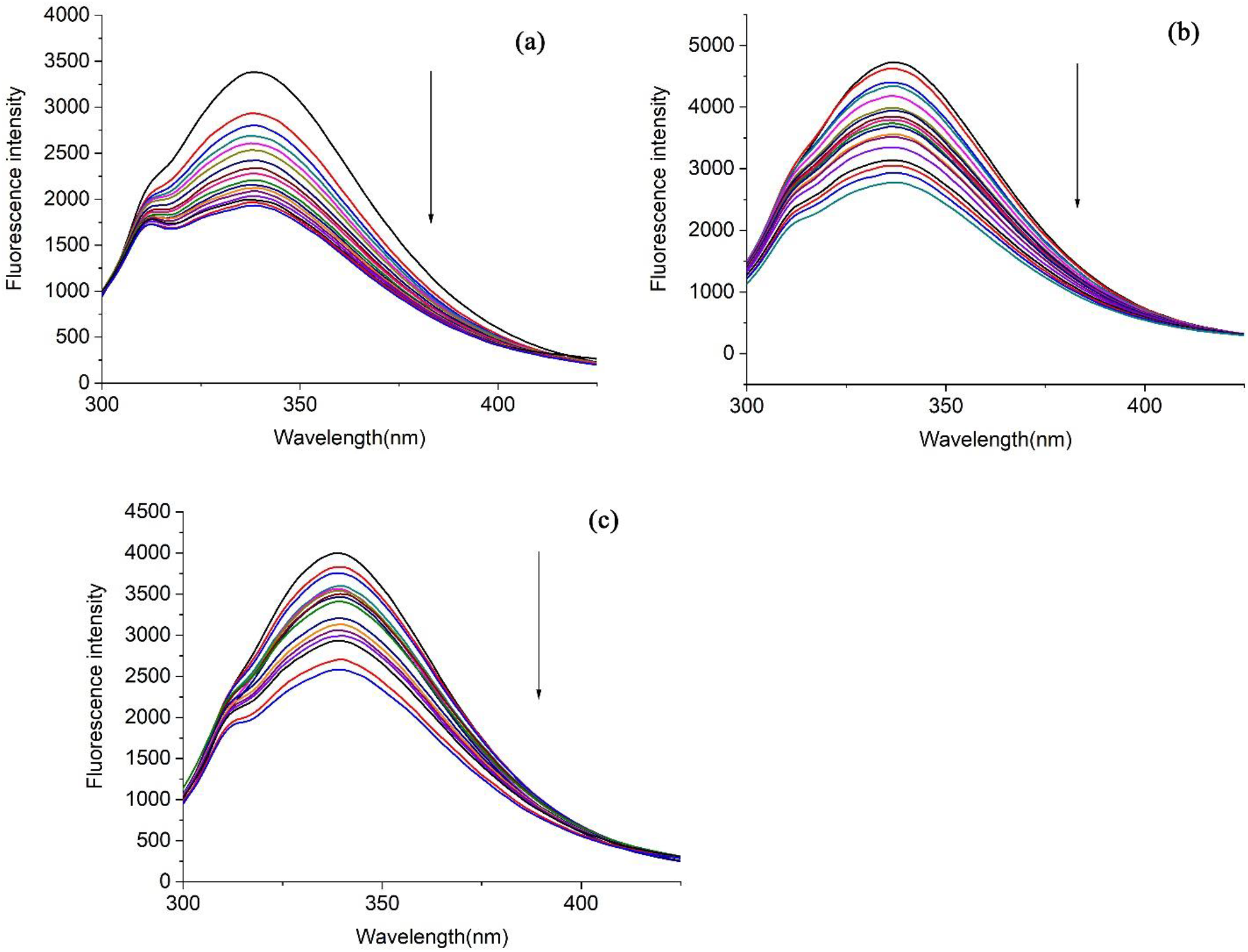
shows the fluorescence spectrum after gradually adding cisplatin (1 μmol/L) to E-cadherin solution (0.2 μmol / L) under different temperature conditions. (a)25°C,(b)31°C(c)37°C

The stern Volmer equation can be used to analyze the fluorescence quenching process [13, 14]

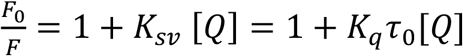

In this formula, F_0_ is the fluorescence intensity of the solution when cisplatin is not added, F is the fluorescence intensity of the cadherin solution after cisplatin is added, [Q] is the concentration of cisplatin, K_SV_ is the quenching constant, and *K*_*q*_ is the quenching rate constant. τ_0_ is the average lifetime of fluorescent molecules in the absence of quenchers.

The quenching constants Ksv calculated using stern Volmer equation under different temperature conditions are listed in Table 1. It can be seen from table 1 that the quenching constant Ksv decreases with the increase of temperature, indicating that the quenching of cadherin by cisplatin could be static quenching. In addition, *K*_*q*_ was calculated from the K_SV_ values is much larger than the maximum dynamic quenching rate constant (2.0× 10^10^ L/ (mol·s) [15], based on the fluorescent lifetime of biomacromolecules is generally in the order of 10^−8^ s [16]. This further reveal that the quenching of cadherin by cisplatin is a static quenching process, and a complex could be formed between cisplatin and cadherin. In the static interaction, the relationship between the quenching fluorescence intensity and the concentration of quencher can be utilized to obtain the expression of the binding constant between the fluorescence quenching molecules. The binding constant (K_b_) and the number of binding sites (n) of cisplatin on cadherin can be analyzed by the following static quenching equation [17].

**Table 1.** Quenching constant and thermodynamic parameters of Cis-Cad(Cis is abbreviated for Cisplatin; Cad is abbreviated for cadherin)

|  | **K<sub>sv</sub>**<br>**(L mol<sup>-1</sup>)** | **K<sub>b</sub>**<br>**(L mol<sup>-1</sup>)** | **n** | **ΔH**<br>**J·mol<sup>-1</sup>** | **ΔS**<br>**J·mol<sup>-1</sup>** | **ΔG**<br>**J·mol<sup>-1</sup>·K<sup>-1</sup>** |
| --- | --- | --- | --- | --- | --- | --- |
| Cis+Cad(25°C) | 2.07×10<sup>7</sup> | 3.20×10<sup>6</sup> | 0.889 |  |  | -1.71×10<sup>4</sup> |
| Cis+Cad(31°C) | 1.66×10<sup>7</sup> | 1.36×10<sup>6</sup> | 0.865 | -3.92×10<sup>4</sup> | -73.96 | -1.67×10<sup>4</sup> |
| Cis+Cad(37°C) | 1.17×10<sup>7</sup> | 8.22×10<sup>5</sup> | 0.86 |  |  | -1.63×10<sup>4</sup> |

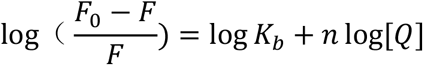

Where F_0_ and F are the fluorescence intensity of the solution without and with cisplatin, respectively, and the intercept is log k_b_. Table 1 lists the calculated values of K_b_ and n under different temperature conditions. It can be seen from Table 1 that K_b_ decreases significantly with the increase of temperature, indicating that the stability of the complex of cadherin and cisplatin decreases significantly with the increase of temperature. The n values at different temperatures are approximate 1, indicating that cisplatin has one binding site on cadherin.

### 3.2 Thermodynamic properties of interactions between cisplatin and cadherin

In the current research system, the temperature change is relatively small, and the enthalpy change ΔH of the binding reaction of cisplatin and cadherin can be approximately considered as a constant.Then the thermodynamic parameters free energy change ΔG, enthalpy change ΔH and entropy change Δ S can be obtained by the following free energy formula and van’t Hoff equation. Table 1 lists the calculated ΔH, Δ S and ΔG in this experiment,.

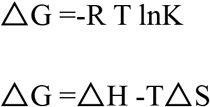

The negative value of ΔG indicates that the binding of cisplatin and cadherin is a spontaneous process. ΔH < 0, indicating that the formation of the complex is an exothermic process. The thermodynamic parameters of the interaction between small molecules and proteins can be used to judge the type of force between them. According to the research on the relationship between thermodynamic data and various forces in the process of binding small molecules to proteins by Ross et al.[18], in the present experiment, ΔH and ΔS <0, indicating that the binding process is dominated by hydrogen bonding and Van der Waals force.

### 3.3 Analysis of atomic force microscope (AFM) imaging results

Atomic force microscopy (AFM) has the advantages for simple sample preparation and intuitive imaging at single molecules level. In recent years, AFM can be used to study proteins and their assembly structures at the single molecule level [19-25]. Analyzing the adsorption behavior and assembly structure of biomacromolecules at the single molecule level can be of great significance for study and cognition of their physiological activities. The high resolution structure and assembly structure of human serum albumin adsorbed on the surface of HOPG have been observed by AFM [22]. In this study, the structures of cadherin and the effect of cisplatin on cadherin assembly were further investigated by AFM.

Fig. 2 shows a typical AFM images of cadherin before and after the interactions with cisplatin at different temperature conditions. It can be seen that the morphology of cadherin changed significantly after the interaction with cisplatin. In Fig. 2 (a), cadherin that does not interact with cisplatin appears as a spherical chain structures and a cross chain structure. After interaction with cisplatin, the length of the chain structure formed between cadherin molecules is significantly reduced, as shown in Fig. 2 (b). In comparing the AFM images under different temperature conditions (Fig. 2 (b), (c) and (d)), the results show that the interaction temperature has a significant effect on the assembly structures of cadherin molecules. With the increasing of the temperature, the chain structure formed between cadherin molecules is further reduced. In Fig. 2 (d), it is judged that there are some cadherin oligomers according to their morphology.

**Fig. 2.**
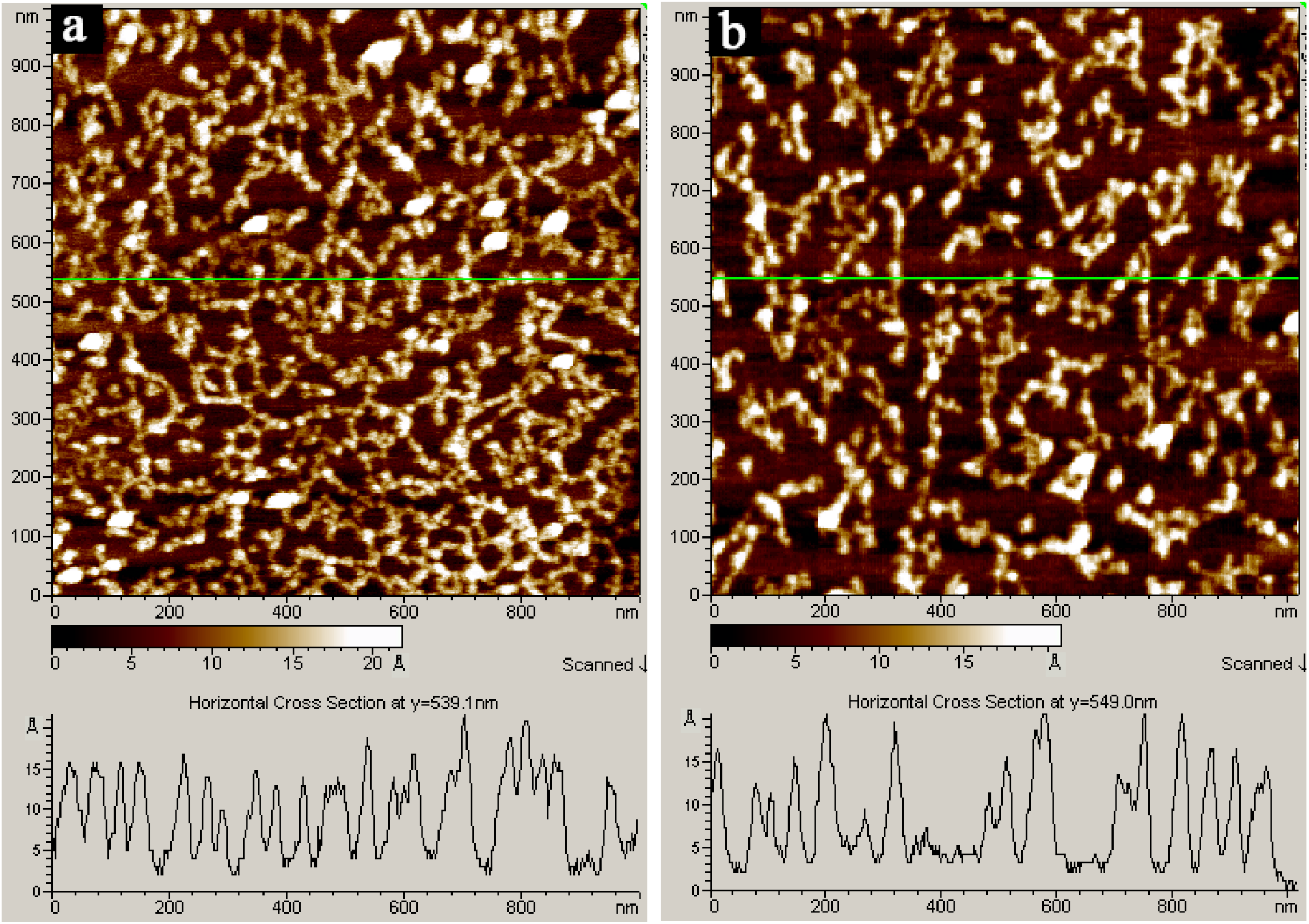
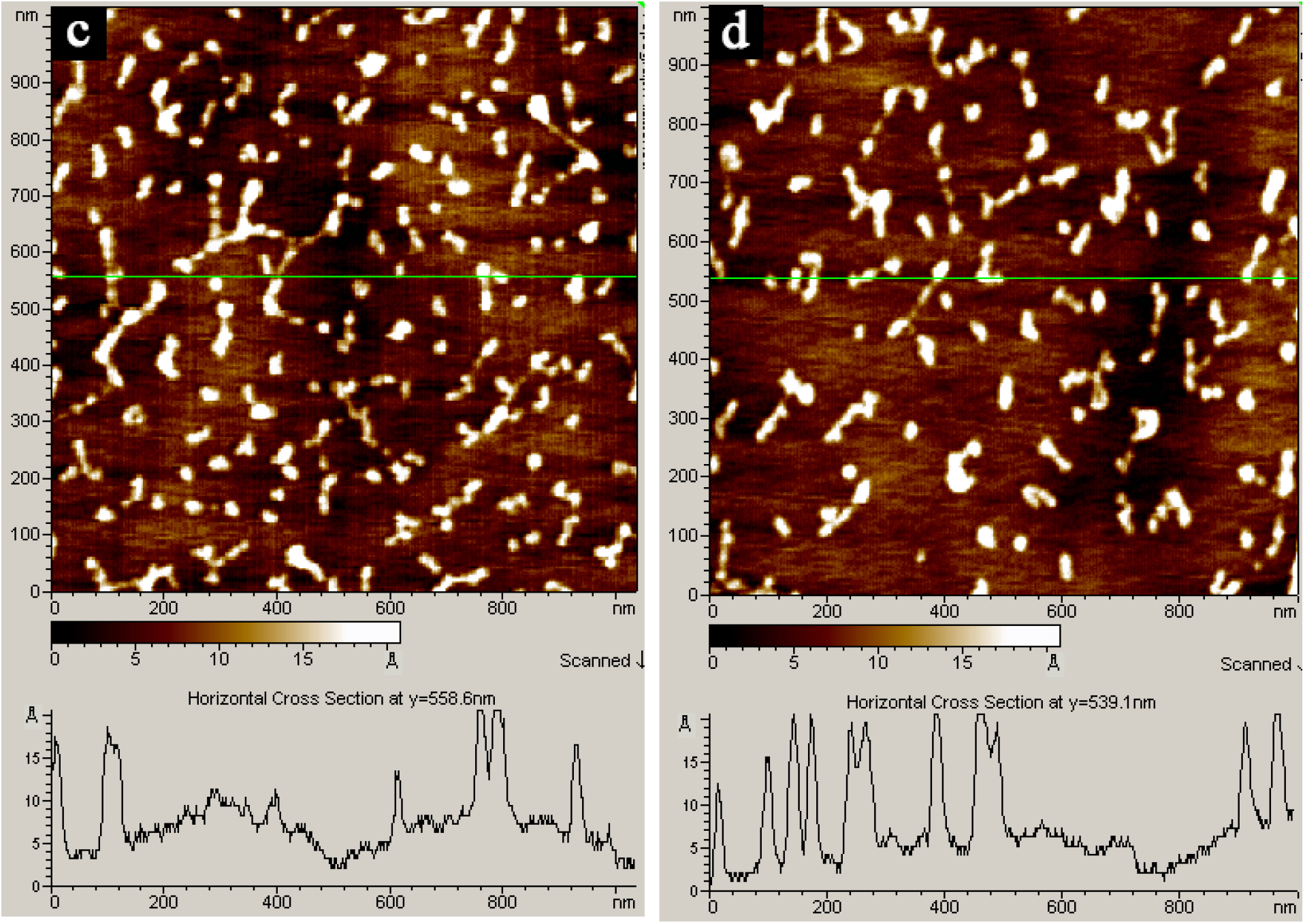
The representative AFM image of 0.02 μM cadherin (a), Cis-Cad at 25°C (b), Cis -Cad at 31°C (c) and Cis -Cad at 37°C (d) on HOPG surface.

The high-resolution structure of cadherin outside the cell membrane has been investigated recently [26-29]. The results of crystal structure study show that each cadherin molecule outside the cell membrane is composed of five repeated domains connected head-to-end(**EC domain 1-5**)[30]. Each domain is a cylindrical structure with a three-dimensional dimension of 4.5 nm × 2.5 nm × 2.5 nm. Because there is a staggered interaction when every two domains are connected, the whole cadherin molecule appears as a chain molecule with a certain arc, and the total length of its head to tail is 20.7 nm. There is a hydrophobic cavity at the N-terminal (EC1 domain) of cadherin molecule, and there is also a tryptophan residue at its distal side. Liquid phase NMR and crystal structure studies show that stable dimers can be formed between two cadherin molecules, in which tryptophan residues of each cadherin molecule can enter the hydrophobic cavity of the other molecule for chain exchange to form a stable structure [30]. The results of high-resolution electron microscopy show that cadherin molecules can also form more complex structures. For example, three cadherin molecules can form a cross structure, and four cadherin molecules can form a four chain structure [26]. In addition to the N-terminal EC1 domain, the force spectrum experimental study shows that there may also be cross domain interaction between cadherin molecules, such as the interaction between EC1 domain and other domains of another molecule [27].

The chain structure and the crossed chain structure observed by the AFM in Fig. 2 (a) can be explained as follows. The spherical chain structure in Fig. 2 (a) may be formed by the interaction between the EC1 domain of the cadherin molecule and other domains of the cadherin molecule due to the cross interaction derived from chain exchange. The cross structure may be formed by the chain exchange of tryptophan residues at the N-terminal (EC1 domain) of three cadherin molecules, which indicates that there is an interdisciplinary interaction between domains among cadherin molecules. In Fig. 2 (b), when the cadherin molecule interacts with cisplatin, the length of the chain structure significantly decreases, mainly because cisplatin interacts with the tryptophan residue at the N-terminal (EC1 domain) of the cadherin molecule, resulting in fluorescence quenching. As confirmed by the fluorescence results, the interaction between the EC1 domain of cadherin and other intermolecular domains of cadherin could be dramatically affected after the cisplatin binding, which is not conducive to the formation of a long chain structure. As the interaction temperature increases to 31 °C and 37 °C, as shown in the AFM images in Fig. 2 (c) and (d), the chain structures further decreases, mainly because the higher temperature is more unfavorable for the cross interaction between the cadherin EC1 domain and other cadherin intermolecular domains. In Fig. 2 (d), it is judged that there are some cadherin oligomers according to their morphology. The average height of the measured cadherin molecule polymer is around of 2.0 nm, which is close to the calculated value of the crystal structure research results (the diameter of cadherin molecule is about 2.5 nm). The formation of cadherin oligomers among molecules is due to the chain exchange of tryptophan in the N-terminal EC1 domain among multiple cadherin molecules, which is consistent with the results of NMR and crystal structure studies on the formation of cadherin oligomers.

## 4 Conclusion

In this study, the interaction between cisplatin and cadherin was analyzed by fluorescence spectroscopy and AFM. The results of fluorescence spectroscopy show that cisplatin can significantly quench the fluorescence emitted by cadherin. It is inferred that this is a static quenching process by calculating the quenching constant and its trend with temperature. The entropy, enthalpy and free energy values of the interaction between cisplatin and cadherin indicate that their binding is a spontaneous process dominated by hydrogen bonds and Van der Waals forces. High resolution AFM showed that long length of the spherical chain structure formed between cadherin molecules. After the interaction with cisplatin, these structures decreased significantly, and became more significant with the increase of temperature. At higher temperature, the formation of a oligomers structure between cadherin molecules was observed. The present AFM results confirmed that there are strong interactions between cisplatin and cadherin, which could affect the activity of cisplatin.

## Acknowledgments

The authors gratefully acknowledge the financial support by the Fundamental Research Funds for the Central Universities (2572020DR07), Natural Science Fund of Heilongjiang Province (LH2019B001), the 111 Project (B20088), Heilongjiang Touyan Innovation Team Program (Tree Genetics and Breeding Innovation Team)

